# Genetic diversity and spread dynamics of SARS-CoV-2 variants present in African populations

**DOI:** 10.1101/2022.08.17.504290

**Authors:** Desire Mtetwa, Tapiwa Fidelity Mutetwa, Tafadzwa Manjengwa, Allen Mazadza, Zedias Chikwambi

## Abstract

The dynamics of COVID-19 disease have been extensively researched in many settings around the world, but little is known about these patterns in Africa. 6139 complete nucleotide genomes from 51 African nations were obtained and analyzed from the National Center for Biotechnology Information (NCBI) and Global Initiative on Sharing Influenza Data (GISAID) databases to examine genetic diversity and spread dynamics of SARS-CoV-2 lineages circulating in Africa. We investigated their diversity using several clade and lineage nomenclature systems, and used maximum parsimony inference methods to recreate their evolutionary divergence and history. According to this study, only 193 of the 2050 Pango lineages discovered worldwide circulated in Africa after two years of the COVID-19 pandemic outbreak, with five different lineages dominating at various points during the outbreak. We identified South Africa, Kenya, and Nigeria as key sources of viral transmissions between Sub-Saharan African nations because they had the most SARS-CoV-2 genomes sampled and sequenced. These results shed light on the evolutionary dynamics of the circulating viral strains in Africa. Genomic surveillance is one of the important techniques in the pandemic preparedness toolbox and to better understand the molecular, evolutionary, epidemiological, and spatiotemporal dynamics of the COVID-19 pandemic in Africa, genomic surveillance activities across the continent must be expanded. The effectiveness of molecular surveillance as a method for tracking pandemics strongly depends on continuous and reliable sampling, speedy virus genome sequencing, and prompt reporting and we have to improve in all these aspects in Africa. Additionally, the pandemic breakout revealed that current land-border regulations aimed at limiting virus’s international transmission are ineffective and a lot needs to be done to implement and improve our African land-borders as far as epidemiology is concerned in order to contain such outbreaks in the future.

## INTRODUCTION

In December 2019, the severe acute respiratory syndrome coronavirus 2 (abbreviated SARS-CoV-2) and the disease caused as a result of the infection is named coronavirus disease (abbreviated COVID-19) was originally identified in Wuhan, China, as a novel virus that was producing a cluster of atypical pneumonia cases [1]. Its outbreak soon became a worldwide pandemic that resulted in a global public health emergency [2]. COVID-19 has affected all seven continents, with Africa being the least stricken by the pandemic [3].

The first confirmed case in Africa occurred on 14 February 2020, in Egypt, while the first case reported in Sub-Saharan Africa occurred on 27 February 2020, in Nigeria [4]. The first cases in other African countries were recorded in March 2020 [5], including in Ghana on 12 March 2020. By 2 May 2021, Africa CDC had reported nearly 4.57 million confirmed cases and more than 122 thousand deaths, as part of the more than 152 million confirmed cases and more than 3.19 million deaths reported globally by WHO SARS-CoV-2 is the third novel coronavirus to be associated with significant outbreaks in the twenty-first century and the seventh coronavirus known to infect people [6]. The first was the SARS-CoV in 2003, which also appeared in China [7], and the second was the Middle East Respiratory Syndrome Coronavirus (MERS-CoV) in 2012, which appeared in Saudi Arabia [7][8]. SARS-CoV-2 is more contagious but appears to have a lower case fatality rate (CFR) than SARS-CoV and MERS-CoV, which both cause serious illness in humans [9][1].

The emergence of SARS-CoV-2 on the African continent necessitates a thorough examination of the virus’s genomic and evolutionary patterns. Comparative analysis of viral genome sequences is an extremely useful method for gaining insight into pathogen emergence and evolution. As a result, this study will conduct an in-depth investigation into the epidemiology and evolution of SARS-CoV-2 in Africa in order to shed light on pandemic dynamics and inform the development and implementation of control measures on the African continent.

## METHODOLOGY

### Dataset mining and workflow

SARS-CoV-2 genome sequences collected from Africa were obtained from the National Center for Biotechnology Information (NCBI) database (https://www.ncbi.nlm.nih.gov) and GISAID database (https://gisaid.org) on July 11, 2022, by only selecting complete genomes excluding those with low coverage using NextClade (https://clades.nextstrain.org). As of 11 July 2022, a total of 3925 SARS-CoV-2 complete genome sequences from 23 African countries were available in the NCBI database and 7828 sequences from 41 African countries were available on GISAID database making a total of 11753 African sequences. The sequences were aligned using the online version of the MAFFT [10] multiple sequence alignment tool (https://mafft.cbrc.jp/alignment/software/closelyrelatedviralgenomes.html), with the Wuhan-Hu-1 (MN 908947.3) as the reference sequence, and 2572 sequences with more than 5.0% ambiguous letters were removed. Duplicates were removed using goalign dedup software and only high quality African complete sequences remained (n=6139).

### Phylogenetic reconstruction

This dataset was subjected to multiple iterations of phylogeny reconstruction using IQ-TREE multicore software version v1.6.12 [11] and nextclade.

### Lineage classification

The web application Phylogenetic Assignment of Named Global Outbreak Lineages (PANGOLin) was used to classify sequences into their lineages. As this nomenclature system is designed to integrate both genetic and geographical information about SARS-CoV-2 dynamics, the goal was to identify the most epidemiologically important lineages of SARS-CoV-2 circulating within the African continent, as well as the lineage dynamics within African regions and across the continent [12].

### Phylogeographic reconstruction

Variants of concern (VOC), variants of interest (VOI) and Variants under monitoring (VUM) were designated based on the World Health Organization framework as of 20 January 2022. We included one lineage, namely A.23.1 and designated it as VOI for the purposes of this analysis. This lineage was included because it demonstrated the continued evolution of African lineages into potentially more transmissible variants. VOI, VOC and VUM that emerged on the African continent were marked. These were A.23.1 (VOI), B.1.351 and B.1.1.529 (VOC), B.1.640 and B.1.525 (VUM). Genome sequences of these five lineages were extracted from NCBI database for phylogeographic reconstruction. A similar approach to that described above (including alignment using online MAFFT) was employed. Phylogeographic reconstruction for all variants circulating in Africa and all VOI, VOC and VUM was conducted using PASTML [13].and maps done using QGIS.

## RESULTS

### Genetic diversity and lineage dynamics in Africa

Of the 11753 complete genomes retrieved from NCBI and GISAID by 11 July 2022, 6139 remained after removing duplicates and sequences with more than 5% ambiguous letters from the dataset. Ancestral location state reconstruction of the dated phylogeny (hereafter referred to as discrete phylogeographic reconstruction) allowed us to infer viral imports and exports between African countries. The early stage of the pandemic was marked by the dominance of lineage B.1, a large European lineage that nearly wiped out Italy (the origin of which roughly corresponds to the Northern Italian outbreak early in 2020). Following the emergence of B.1.351 (Beta) in South Africa, it became the most frequently detected SARS-CoV-2 lineage in Africa during the pandemic’s second wave. Due to the relaxation of travel restrictions, other variants of concern, such as the B.1.1.7 (Alpha) UK lineage and the B.1.617 (Delta) Indian lineage, made their way into Africa and became dominant during other waves of the pandemic. B.1.525, AY.16, C.36, AY.116, A.23.1, and AY.122 are some notable lineages that dominated during different waves of the pandemic. BA.1, an alias of B.1.1.529.1, also known as Omicron, preceded the fourth pandemic wave and was first reported in South Africa. Egypt reported the first SARS-CoV-2 case in Africa, which was of lineage B. This lineage was exported from other continents, primarily Europe, to other African countries.

### Emergence and spread of new SARS-CoV-2 variants

In order to determine how some of the key SARS-CoV-2 variants are spreading within Africa, we performed phylogeographic analyses on the VOC B.1.351 and B.1.1.529, VUM B.1.525 and B.1.640 and one additional variant that emerged and that we designated as VOI for this analysis A.23.1. Molecular clock analysis of these five datasets provided strong evidence that these five lineages are evolving in a clocklike manner (Fig. 4a – Fig 4e).

**Fig 1:**
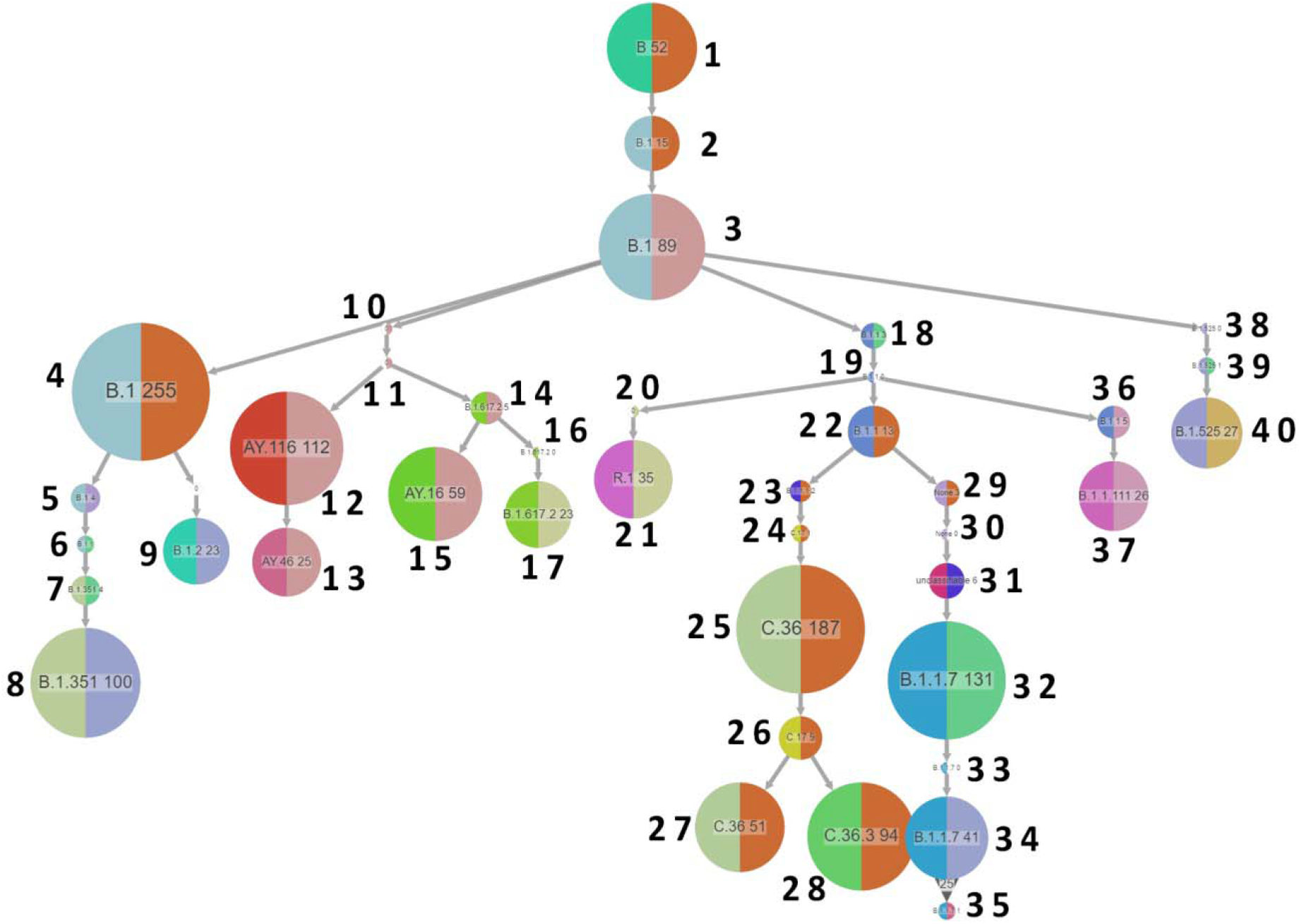
Phylogeographic reconstruction of African countries showing that after entering Egypt, the lineage evolved into the B.1 lineage before spreading to Kenya, Morocco, and Ghana. From there, there were imports and exports of the B.1 lineage between African countries, and the lineage evolved from B.1 into many variants (diversification events) across the continent. Lineage A also entered the picture from other continents, evolved within the continent, and spread across the continent

**Fig 2:**
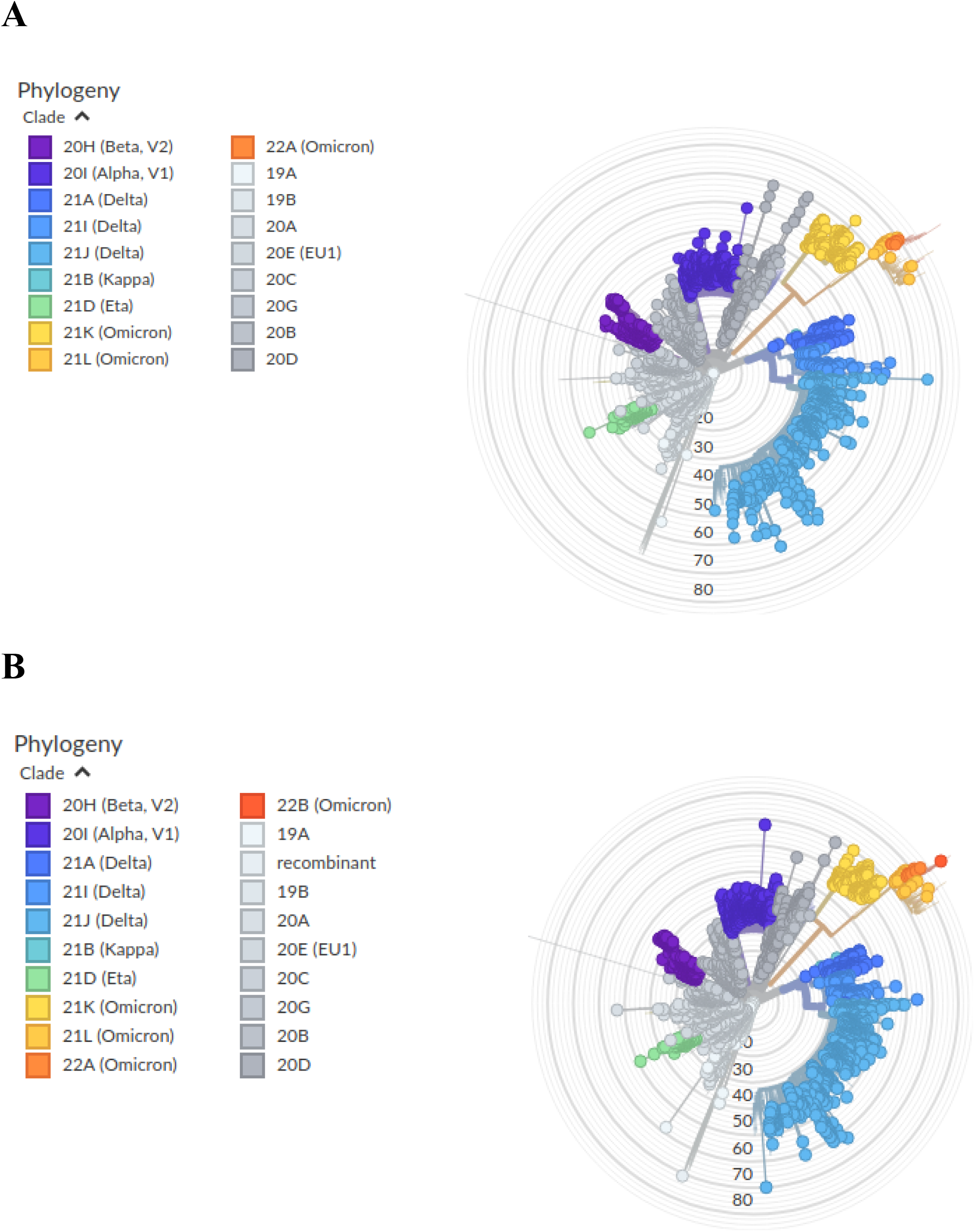
Phylogenetic tree of clades circulating in Africa from 6 139 sequences. A (3 070 sequences) and B (3 069 sequences) showing clades and their number of mutations.

**Fig 3:**
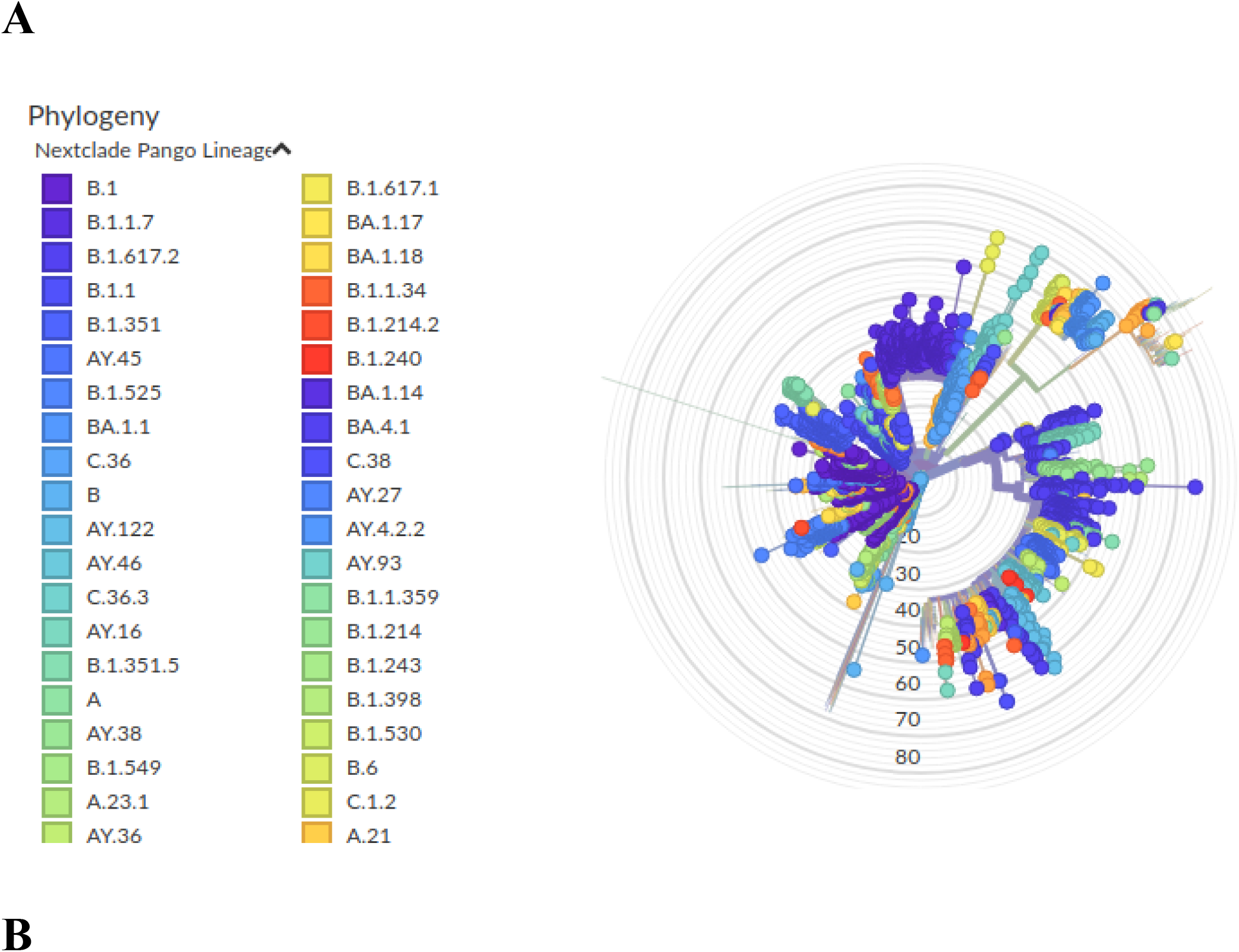
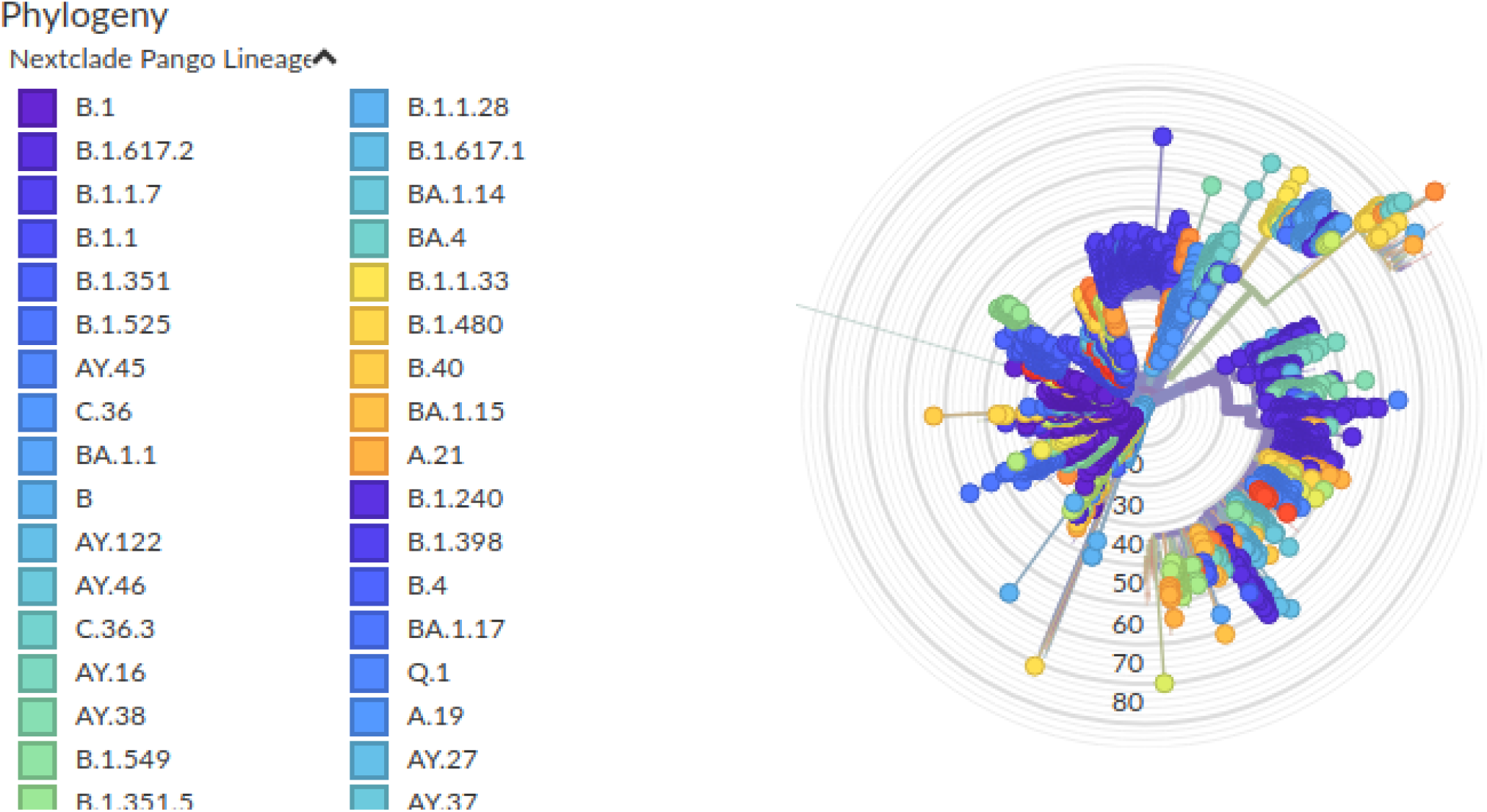
NextClade Phylogenetic tree of lineages circulating in Africa from 6 139 sequences. A (3 070 sequences) and B (3 069 sequences) showing number of mutations,

**Fig 4:**
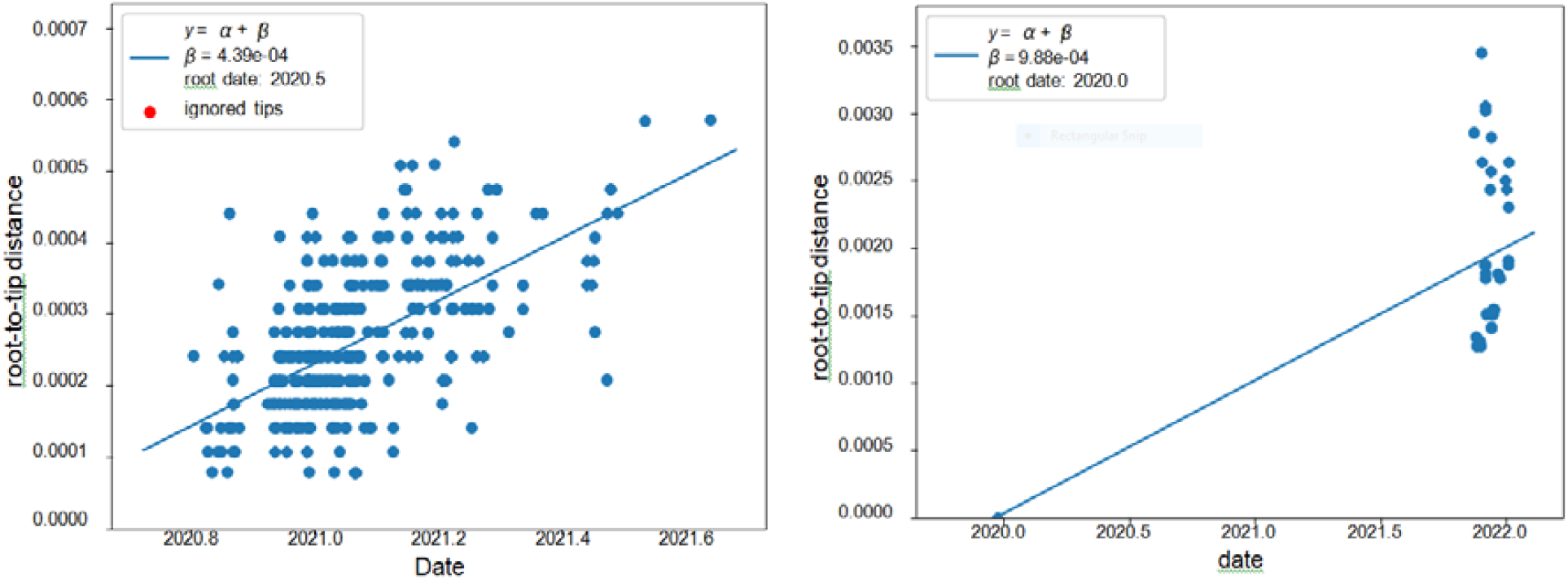
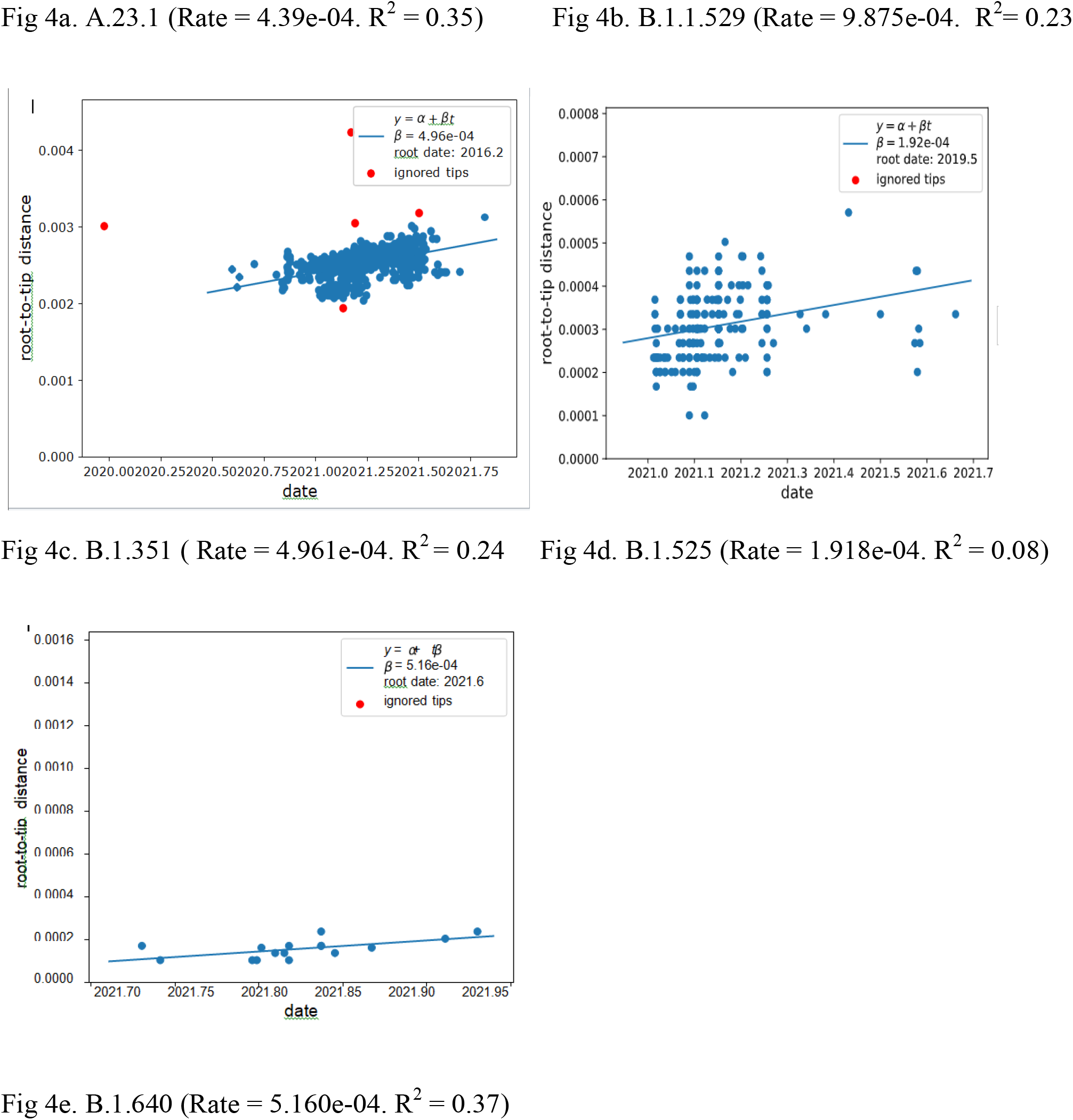
Molecular clock analysis of the Linear-regression curve and co-relationship of the Root to tip distance against time for SARS-CoV 2. Each circle corresponds to a tip in the phylogenetic tree. The y-axis corresponds to the root-to-tip distance of the phylogenetic tree with branch lengths in units of substitutions per site. The x-axis represents calendar time. Adjusted R^2^ is from the linear regression model, beta coefficient of the mobility indicator from the linear regression equation, *p*-value from the linear regression model. The regression line is the best fitting line using the root position that maximized R^2^

### Variants of Concern

#### B.1.351

In October 2020, B.1.351 was sampled for the first time in South Africa. It was defined by ten spike protein mutations.

#### B.1.1.529

South Africa notified the World Health Organization (WHO) on 24 November 2021 that a new SARS-CoV-2 variant, B.1.1.529, had been discovered. B.1.1.529 was initially discovered in specimens taken in Botswana on 11 November 2021, and in South Africa on 14 November 2021. Since then, South Africa has discovered B.1.1.529 in samples taken on 8 November 2021. At least 30 amino acid changes, three modest deletions, and one small insertion characterize the Omicron variant’s spike protein.

### Variants Under Monitoring

#### B.1.525

B.1.525 is a VOI defined by six substitutions in the spike protein, and two deletions in the N-terminal domain

#### B.1.640

B.1.640 is a variant that originated from Central Africa. According to our phylogeographic reconstruction, it was first reported in Congo and it was directly transmitted to Ghana and Kenya It has two sublineages, B.1.640.1 and B.1.640.2, and the B.1.640.2 was a VOC as of February 17, 2022, due to 46 mutations and 37 deletions in its genetic sequence, many of which impair the spike protein.

### Variant of Interest

#### A.23.1

We designated A.23.1 as VOI for the purposes of this analysis, as it present good examples of the continued evolution of the virus within Africa.

## DISCUSSION AND CONCLUSION

The first SARS-CoV-2 case in Africa was first reported by Egypt, followed by Nigeria, then South Africa and then Ghana and Kenya on the same day (12 March 2020). Egypt, Kenya, and South Africa appear to be key suppliers of importations into other African countries, [14] found out that this is likely impacted by the fact that these three nations have the most deposited sequences and are among the first to report cases of SARS-CoV-2 in Africa. Lineage B was the first SARS-CoV-2 case reported in Egypt, indicating that the early introduction in Africa was an Asian lineage reported by China. The early stage of the pandemic was distinguished by the dominance of lineage B. 1. a large European lineage (the origin of which roughly corresponds to the Northern Italian outbreak early in 2020) rather than lineage B, which caused the pandemic’s establishment in China. According to [14] lineage B.1 was introduced multiple times to African countries in the early phase of the pandemic, thus most of the introductions where predominantly from Europe than Asia

Our phylogeographic analysis of African countries reveals that after entering Egypt, the lineage developed into the B.1 lineage before extending to Kenya, Morocco, and Ghana. Following that, the B.1 lineage was imported and exported across African countries, and the lineage diversified into numerous variants (diversification events) across the continent. Lineage A also entered the picture from other continents as air travel restrictions were relaxed, allowing for intra-country and occasional international viral movements between neighboring countries, presumably via road and rail ties. Though some border posts between countries were closed during the initial lockdown period, others remained open to allow trade to continue, and as a result, variants evolved within the continent and spread across it.

When lineage B.1.351 (Beta) first appeared in South Africa, it quickly spread throughout the country because the variant was thought to propagate quicker than previous virus versions. By March 2021, B.1.351 has surpassed B.1. as the dominant lineage in most Southern African and East African countries. During the second wave of the epidemic, it became the most often discovered SARS-CoV-2 lineage in Africa. This occurred owing to the loosening of air and land travel limitations, which also resulted in the arrival of various variants and lineages from other continents. Variants of concern like B.1.1.7 (Alpha) of the UK lineage and B.1.617 (Delta) of the Indian lineage made their way to Africa and became prominent during other pandemic waves. B.1.351 moved directly from Southern Africa to East and Central Africa, according to our phylogeographic reconstruction. A discrete phylogeographic examination of a larger sample of viruses by [14] implies that transmission to West Africa may have occurred via East Africa, with a probable European intermediary.

BA.1, an alias of B.1.1.529.1 also known as Omicron, which was initially detected in South Africa, preceded the fourth pandemic wave. South Africa notified the World Health Organization on November 24, 2021, that a new SARS-CoV-2 variant, B.1.1.529, had been discovered. B.1.1.529 was initially discovered in specimens taken in Botswana on 11 November 2021, and in South Africa on 14 November 2021. Since then, South Africa has discovered B.1.1.529 in samples taken on 8 November 2021.The spike protein of the Omicron variant is characterized by at least 30 amino acid substitutions, three small deletions, and one small insertion. As a result, Omicron was designated as a VOC. B.1.1.529 extended into Botswana and South Africa’s neighboring countries, then into Northern and Western Africa (Morocco and Nigeria), as well as into their neighboring countries. (Fig 5b). Other transmissions from Botswana to Mali and Ghana have been documented. Clearly, the variety was reintroduced to the Southern countries and other countries in the time between transmissions. As a result, most Southern African countries were placed on the blacklist, and travel to other countries was prohibited.

**Fig 5a:**
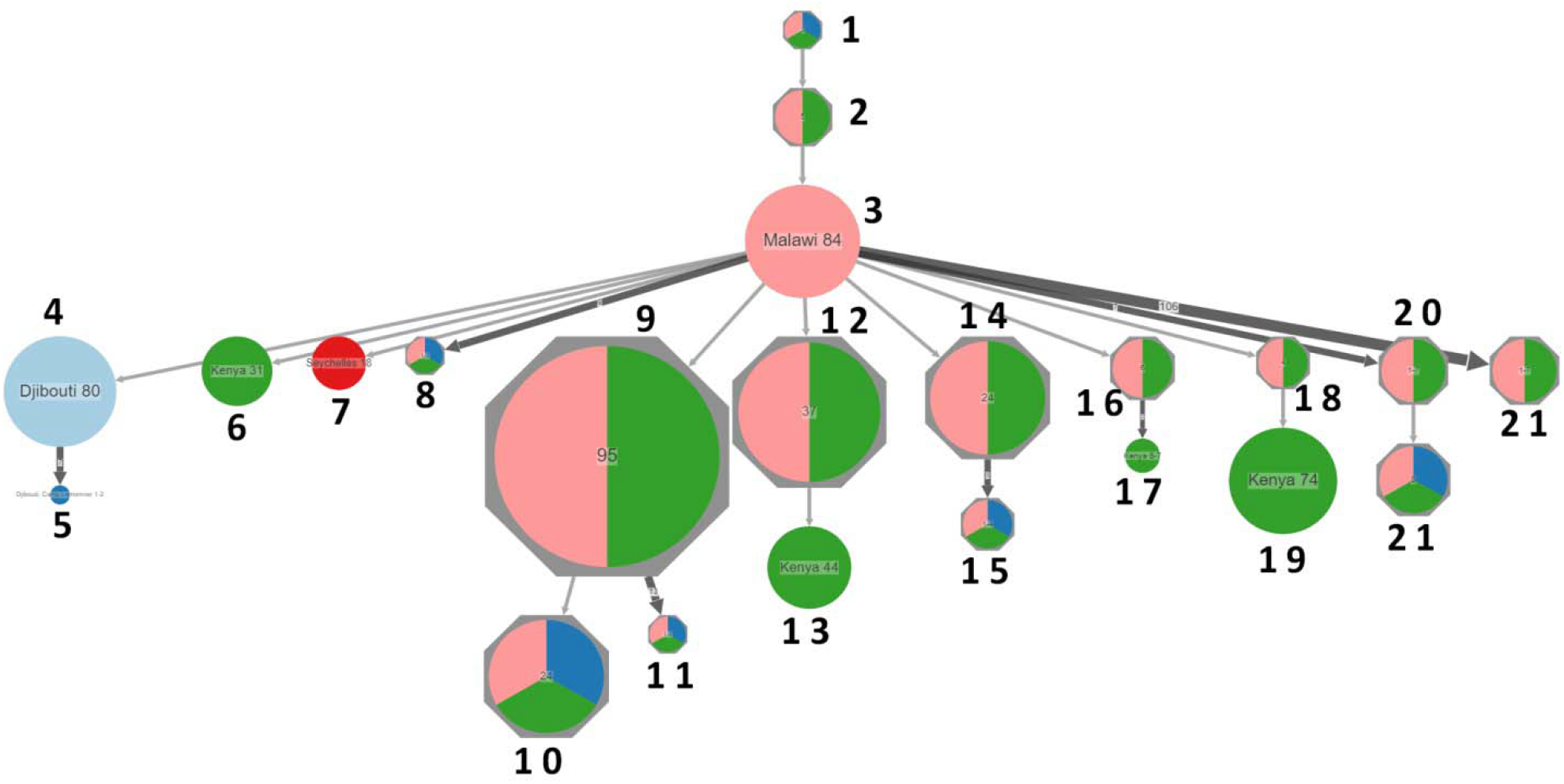
Spread dynamics of B.1.351, It was first sampled in South Africa. Following its emergence in the Eastern Cape, the virus spread rapidly throughout South Africa, as the variant spreads faster than previous variants of the virus. The variant was responsible for the pandemic’s second wave in South Africa. The variant had spread into neighboring countries by November 2020, and by December 2020, it had reached Djibouti and Malawi. By March 2021, B.1.351 had become the dominant lineage in most Southern African and East African countries. B.1.351 also moved directly from Southern Africa to East and Central Africa.

**Fig 5b:**
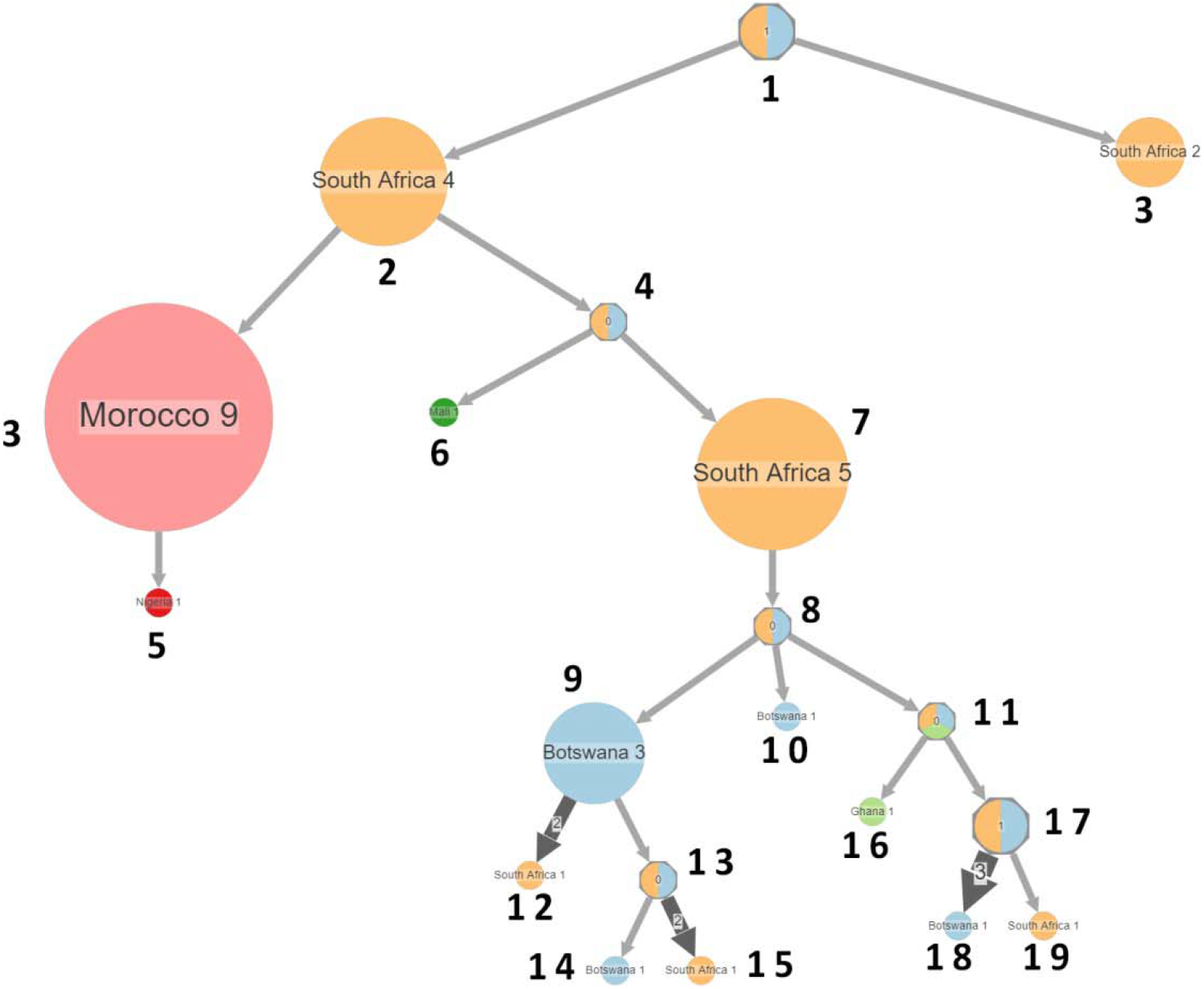
Spread dynamics of VOC B.1.1.529 expanded into Botswana and South Africa’s neighboring countries, then into Northern and Western Africa (Morocco and Nigeria), as well as into their neighboring countries. Other transmissions from Botswana to Mali and Ghana have been documented. Obviously in-between transmissions, re-introduction of the variant to the Southern countries occurred..

**Fig 5c:**
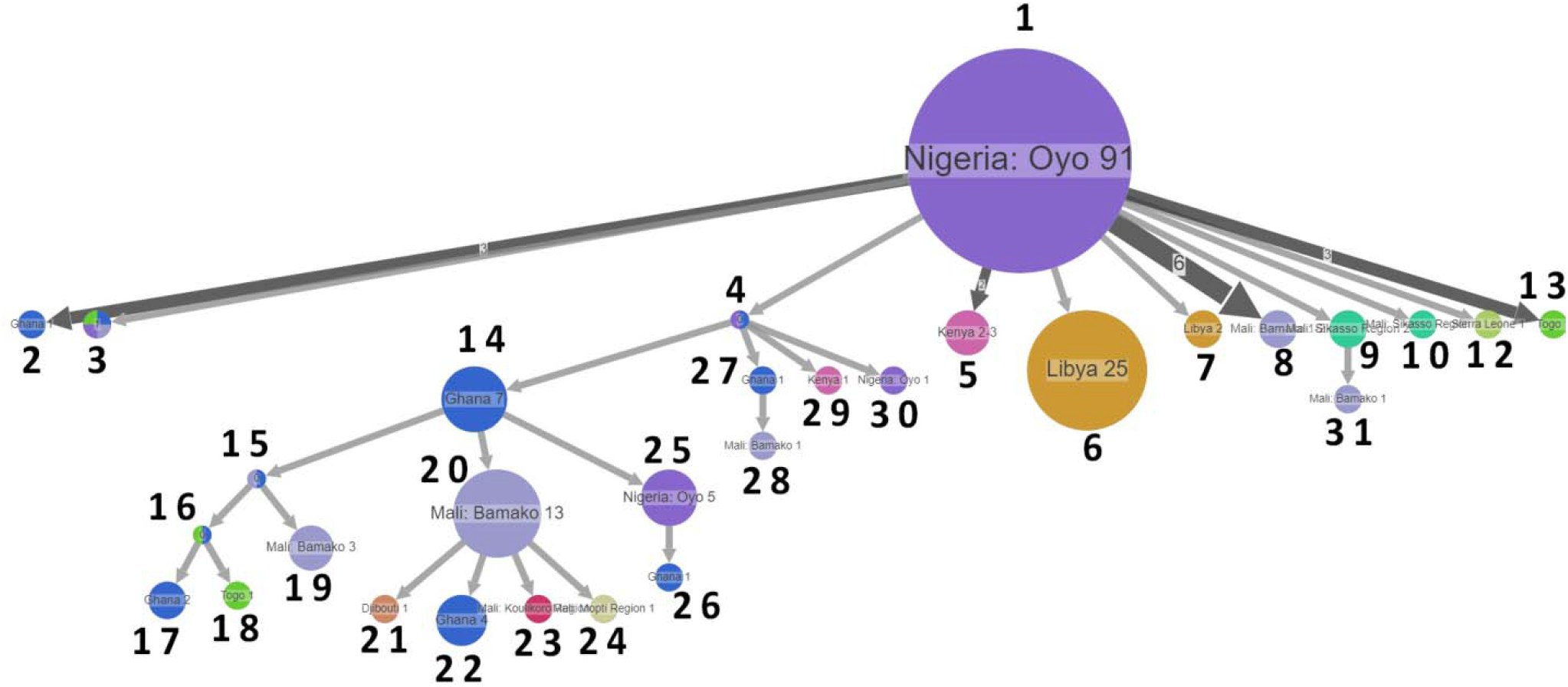
Spread dynamics of B.1.525 demonstrates movement of B.1.525 into neighboring countries like Ghana, Mali, Togo, Sierra Leone, East Africa like Kenya, Djibouti and Northern Africa like Libya directly from Nigeria

**Fig 5d:**
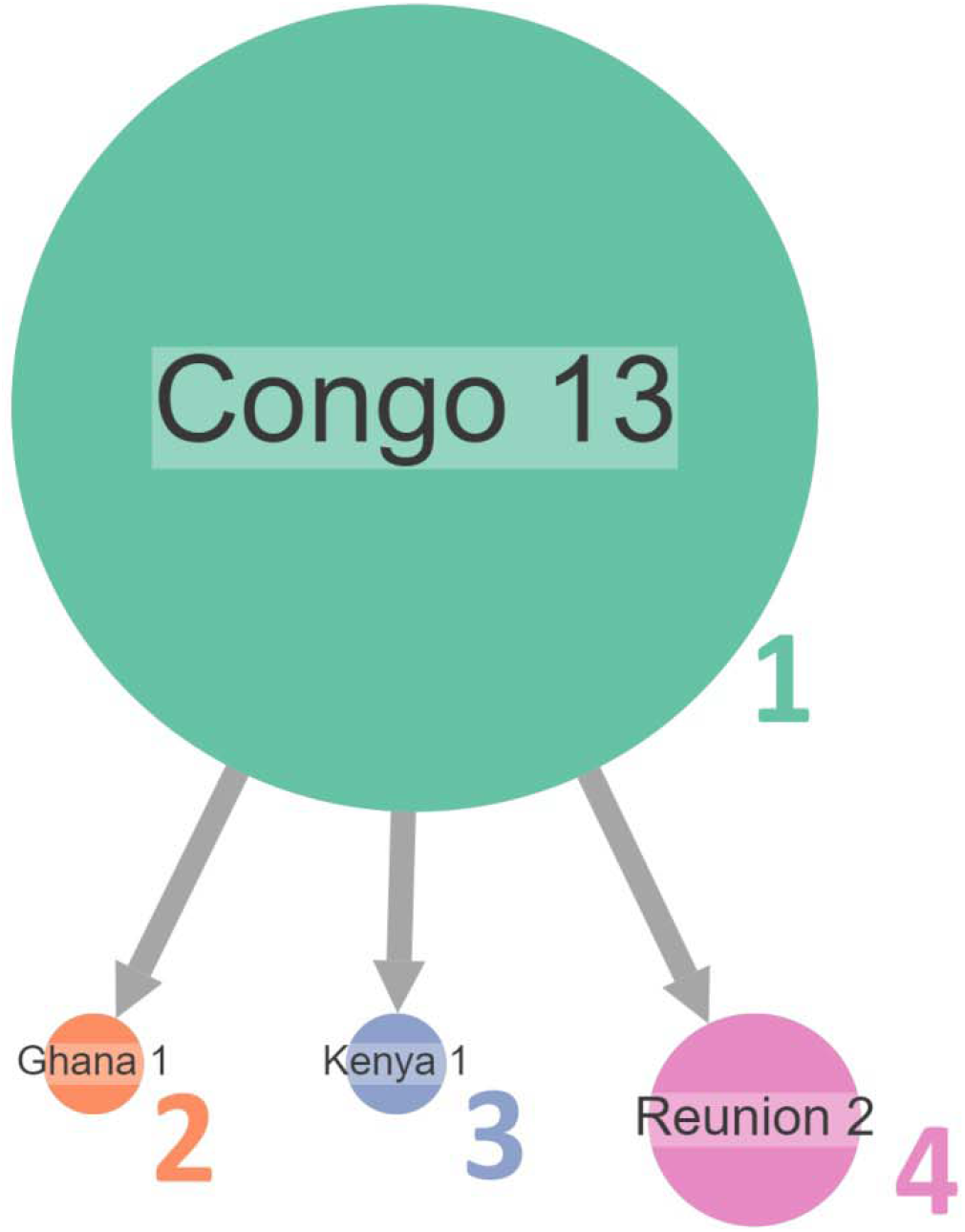
Spread dynamics of B.1.640. It originated from Central Africa and it was first reported in Congo. It was directly transmitted to Ghana and Kenya

**Fig 5e:**
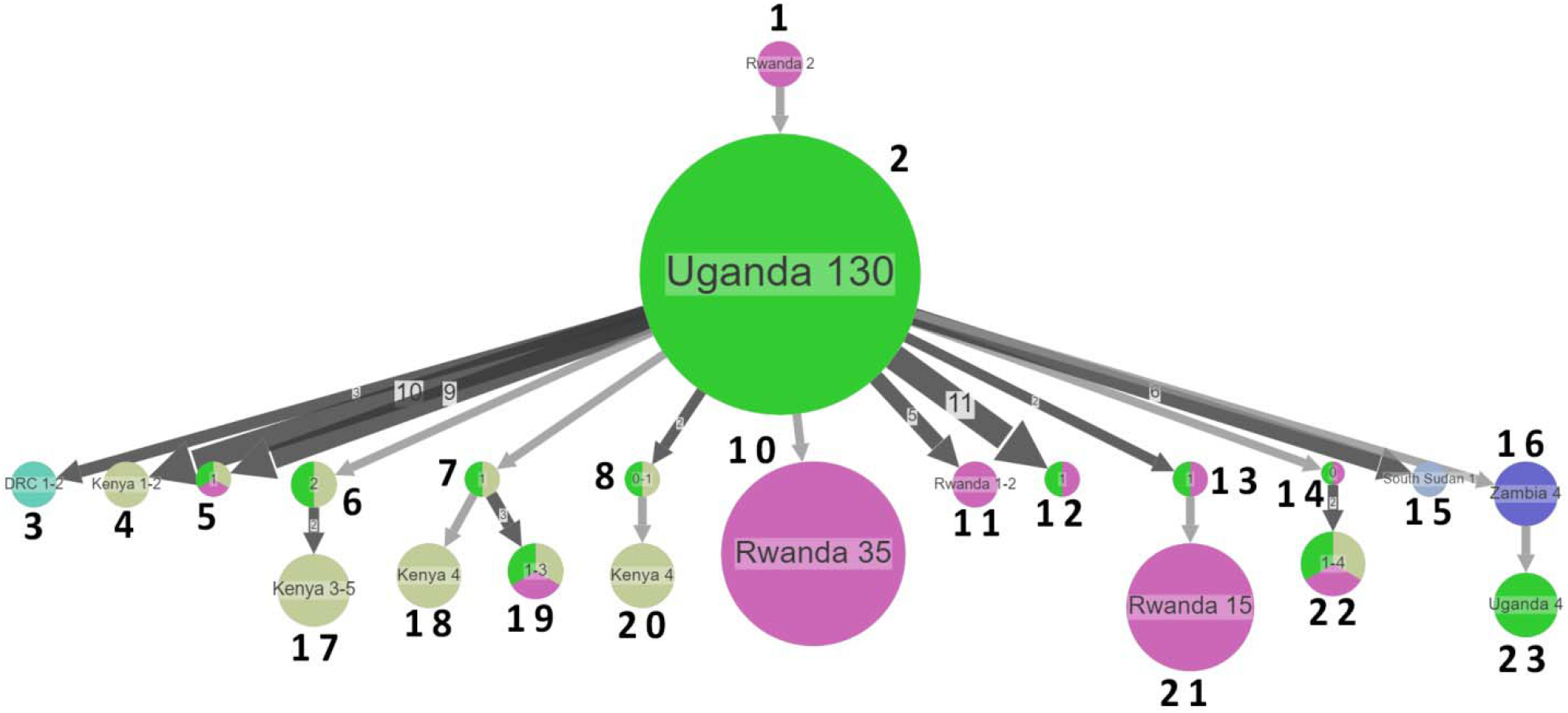
Spread dynamics of A.23.1. Since its discovery in September 2020, A.23.1 has spread regionally into neighboring Rwanda and Kenya, as well as to the DRC, South Sudan, and Zambia in the south.

**Fig 6.**
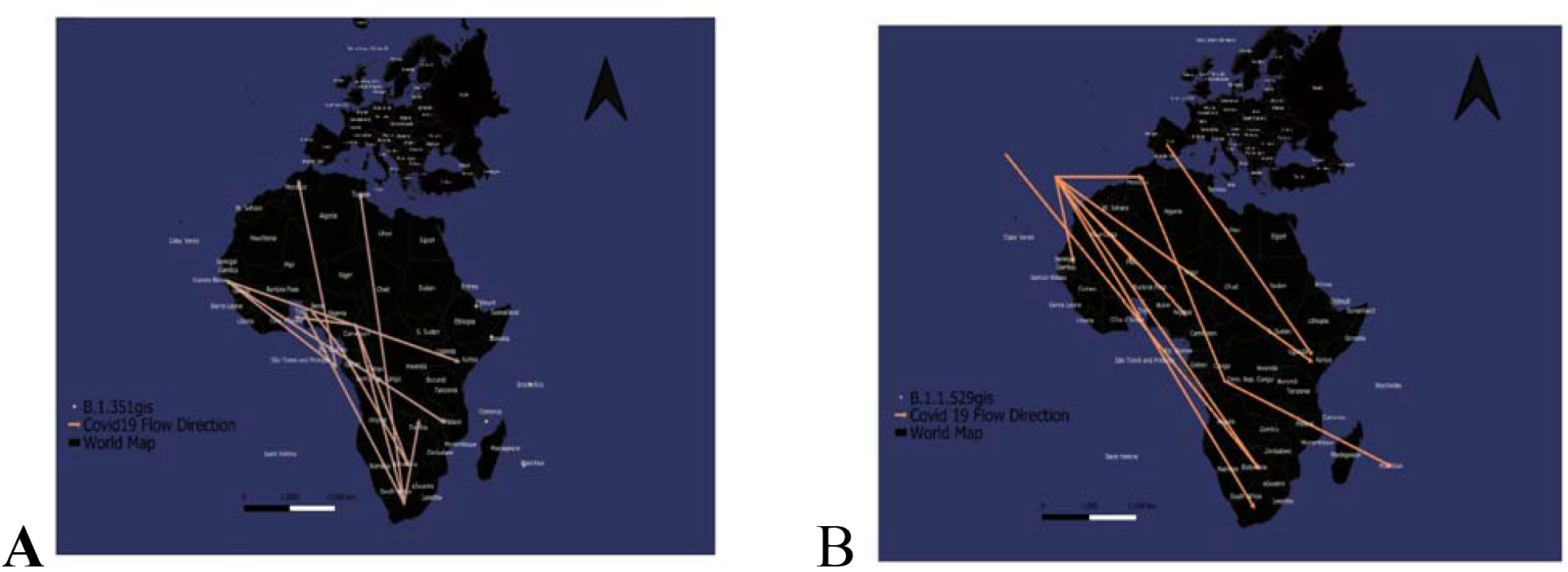
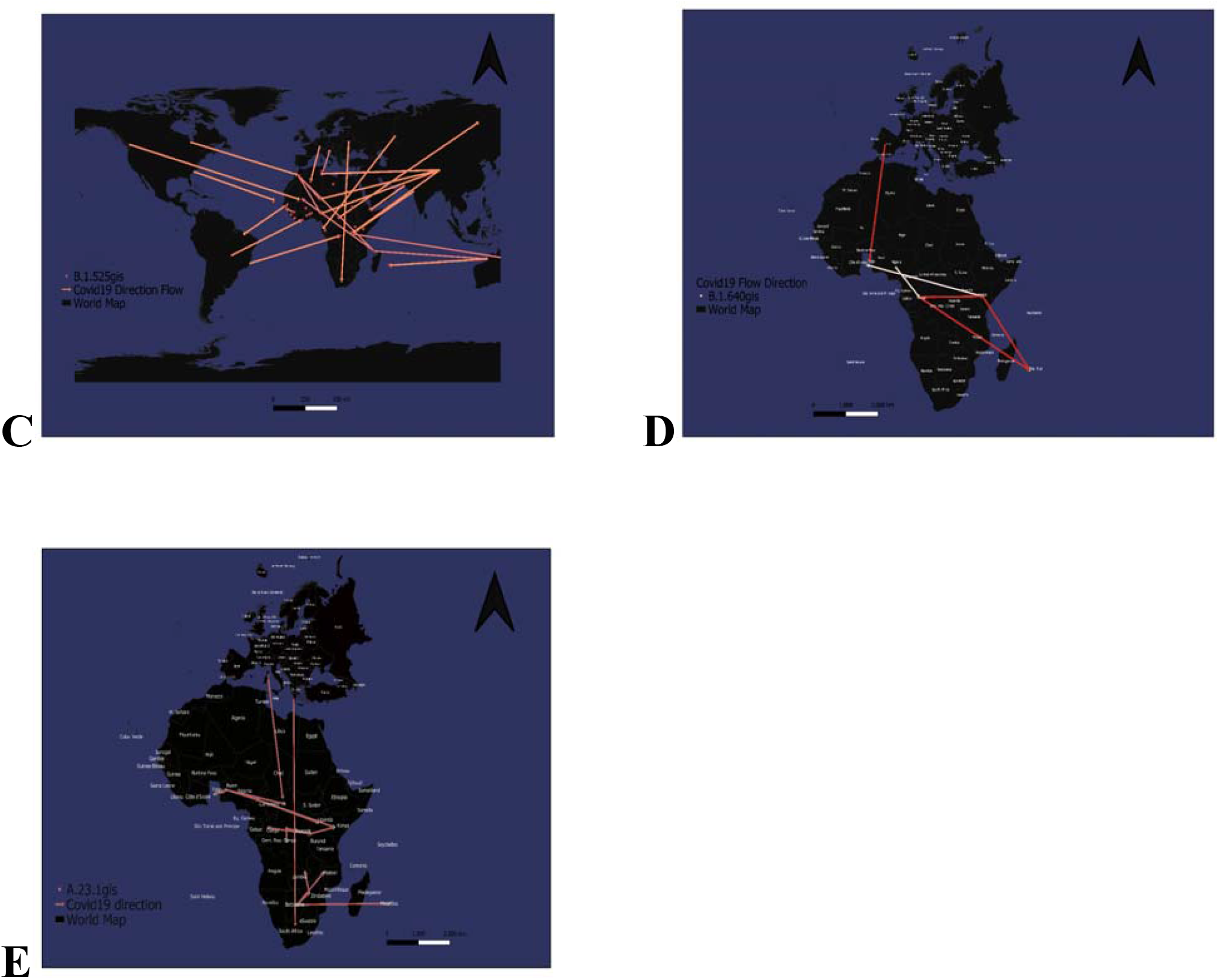
Spread dynamics of A (B.1.351) B (B.1.1.529) C (B.1.525) D (B.1.640) and E (A.23.1) within African continent

B.1.525 is a VOI defined by six substitutions in the spike protein and two deletions in the N-terminal domain. According to [14] this was first sampled in the United Kingdom in mid-December 2020, but their phylogeographic reconstruction suggested that the variant originated in Nigeria in November 2020. Since then it has spread throughout much of Nigeria and neighbouring Ghana. Our phylogeographic reconstruction also demonstrates movement of B.1.525 into neighboring countries like Ghana, Mali, Togo, Sierra Leone, East Africa like Kenya, Djibouti and Northern Africa like Libya directly from Nigeria (Fig 5c). The scope of this VOI’s distribution in the region is unknown due to poor sampling from other neighboring nations in West and Central Africa.

B.1.640 is a variant that originated from Central Africa. According to our phylogeographic reconstruction, it was first reported in Congo and it was directly transmitted to Ghana, Kenya and Reunion (Fig 5d). Given the small sample size from all African countries, the extent of this VUM’s spread in the region is unknown. What we know is that it has two sublineages B.1.640.1 and B.1.640.2 and as of 17 February 2022 the B.1.640.2 was a VOC because of its 46 mutations and 37 deletions in its genetic code, many affecting the spike protein. This sublineage also dubbed as the IHU was on the rise in France.

We designated A.23.1 as VOI as it presents good examples of the continued evolution of the virus within Africa. According to [15], the variant contains several mutations observed in variants of concern as well as six unique substitutions. However, the variant does not have a similar origin with all of the variants of interest or concern, including alpha, beta, gamma, delta, and mu, [15]. All of those variants share a mutation that identifies their common ancestry.

A.23.1, on the other hand, does not. It has more in common with the A.30 variety, which was first discovered in Angola and may have originated in Tanzania. According to [15], neither of these variants has a common ancestor with the other major strains, and the identification of two separate yet distantly related variations in East Africa is troubling in and of itself. The fact that these variations originated separately from all others in the globe, despite the fact that they lack the characteristic trio of mutations that link all other current variants, illustrates the adaptability of SARS-CoV-2 to local conditions. Lineage A.23.1, characterized by three spike mutations (F157L, V367F and Q613H), it was first detected in July 2020, in a Ugandan prison in Amuru. The lineage was then transmitted to Kitgum prison, possibly aided by the transfer of prisoners. Following that, the A.23 lineage infiltrated the general population and spread to Kampala, introducing new spike mutations (R102I, L141F, E484K, P681R) as well as additional mutations in nsp3, nsp6, ORF8 and ORF9, prompting the creation of a new lineage classification, A.23.1 [14]. Since its discovery in September 2020, A.23.1 has spread regionally into neighboring Rwanda and Kenya, as well as to the DRC, South Sudan, and Zambia in the south (Fig 5e). However, according to [14] their phylogeographic reconstruction of A.23.1 suggests that the introduction into Ghana may have occurred via Europe, whereas the introductions into southern Africa likely occurred directly from East Africa. This is consistent with epidemiological data indicating that the case discovered in South Africa was a contact of a person who had recently visited Kenya.

High rates of COVID-19 testing and consistent genomic surveillance in the continent’s south have resulted in the early detection of VOCs like B.1.351 and B.1.1.529. Since the discovery of these southern African variants, several other SARS-CoV-2 VOIs have emerged around the world, including on the African continent, including B.1.525 in West Africa and A.23.1 in East Africa. There is strong evidence that both of these VOIs are becoming more common in the regions where they have been identified, implying that they may be more fit than other variants in these regions.

In the latter months of 2020, this variant moved from South Africa into neighboring nations, reaching as far north as the Democratic Republic of the Congo (DRC) by February 2021, according to our focused phylogenetic analysis of the B.1.351 branch. This spread could have been aided by rail and road networks that connect South Africa’s ocean ports to economic and industrial centers in Botswana, Zimbabwe, Zambia, and the DRC’s southern regions. The virus’s rapid and seemingly unhindered penetration into these countries implies that present land-border regulations aimed at limiting the virus’s international transmission are ineffective and a lot needs to be done to implement and improve our African land-borders as far as epidemiology is concerned in order to contain such outbreaks in the future

From the standpoint of public health, genetic surveillance is just one instrument in the pandemic preparedness toolset. On the other hand, in Africa, genetic surveillance is not being carried out effectively. The usefulness of molecular surveillance as a technique for monitoring pandemics is heavily reliant on ongoing and consistent sampling, quick virus genome sequencing, and timely reporting.

When this is accomplished, molecular surveillance will be able to detect shifting pandemic traits early. Molecular monitoring data can also inform public health interventions when such changes are found. In this context, the molecular surveillance data collected by the majority of African countries is less useful than it could be. For example, in some circumstances, the time lag between when virus samples are acquired and when sequences for these samples are stored in sequence repositories is so great that the genomic surveillance data’s primary utility is lost. Depending on the laboratory or country, numerous variables contribute to this lag: (i) a paucity of reagents due to global supply chain disruptions, (ii) a lack of equipment and infrastructure within the source country, (iii) a scarcity of technical skills in laboratory methods or bioinformatics support, and (iv) a reluctance to share data by some health officials. To disclose the genetic properties of currently circulating viruses in these nations, more recent sampling and fast reporting is required.

As a result, our study’s fundamental flaw is the patchiness of African genetic monitoring data. It’s worth noting that phylogeographic reconstruction of viral propagation is strongly reliant on sampling, with the caveat that the exact paths of viral migrations between nations cannot be deduced if connecting countries are not sampled. Furthermore, uneven sampling between African countries is very certainly biasing our efforts to recreate SARS-CoV-2 migration dynamics across the continent. It’s no surprise that we identified South Africa, Kenya, and Nigeria as key sources of viral transmissions between Sub-Saharan African nations because they had the most SARS-CoV-2 genomes sampled and sequenced.

The strength with which national public health programs embrace genomic surveillance as a tool for preventing the appearance and spread of dangerous variants determines its reliability as a strategy for preventing the emergence and spread of dangerous variants. [14] states that as in most other parts of the world, the success of genomic surveillance in Africa requires that more samples be tested for COVID-19, that a higher proportion of positive samples be sequenced within days of sampling.

The degree of the dissemination of specific variations in the region is unknown due to limited sampling across all African countries. Some variants have been sequenced to exhaustion and deposited in databases, whilst others, such as B.1.640 and B.1.525, have relatively few sequences deposited in databases.

Some countries have a considerable number of sequences deposited in sequence repositories, whereas others have few or none. Our research became skewed as a result of this. PASTML, which we used to do a phylogeographic reconstruction, also required caution with sampling bias; it was necessary to subsample sequences to better represent the number of declared cases by location and/or use additional data, such as travel history, or the phylogeographic predictions would be incorrect. complete genome sequencing of SARS-CoV2 is important to find useful information about the viral lineages, variants of interests and variants of concern, but only about 32% of the nucleotide sequences deposited in NCBI database from Africa are complete with 68% partial nucleotide sequences. As a continent we need to be able to deposit complete genomic sequences because most researchers only take complete genomes for their studies and perhaps because of this, we discovered there are fewer lineages circulating throughout Africa than there actually are.

Since the pandemic began multiple consortiums and collaborations between public and private sectors have been established to sequence and publish the SARS-CoV-2 genomes in real-time. As of 19 May 2022, there were 10,900,329 genome sequences available at GISAID which is the largest number of whole-genome sequencing (WGS) carried out for any organism today. Even though there is an increased number of WGS that are being carried out, we need universal agreement as there is no universal agreement between different consortia as to what are the quality criteria that have to be followed when sequencing or using the publicly available SARS-CoV-2 genomes for analysis. Currently, there are several sequencing strategies, with multiple protocols that are being used that are conserved for that consortium or the country as a whole. Public health laboratories follow their own sample selection criteria, library preparation and sequencing platforms, bioinformatics workflows, and data interpretation, resulting in inconsistent data quality standards and biases among SARS-CoV-2 sequences generated in public databases.

Phylodynamic analyses of SARS-CoV-2 variants showed notably low Rt values which may be due to effective lockdowns, social distancing and mask wearing polices that contributed to the control of SARS-CoV-2 in the African countries indicated. The estimates of the Maximum likelihood phylogenetic relationships among the SARS-CoV-2 genomes for our dataset using the tip-dating method show low genetic diversity. We also inferred a weak yet significant temporal signal in the dataset, that reflects a low mutation rate of SARS-CoV-2, which is consistent with findings reported elsewhere.

## Conclusions

Our findings suggest that SARS-CoV-2 variants circulating in Africa follow unique patterns of evolution and transmission dynamics. Routine genomic surveillance of the variants employing the use of phylogenetic and phylodynamic analyses with a wider coverage and higher resolution is essential for providing insight into the effectiveness of different intervention strategies against the pandemic.

## Author Contributions

DM designed the study. DM participated in the data collection and curation. DM, AM, TapM and TafM performed the analyses. DM, AM, and ZC drafted the initial manuscript. DM, AM and ZC administered and supervised the study. All authors read and approved the final manuscript.

## Acknowledgments

We thank all those who have contributed sequences to the GISAID database (https://www.gisaid.org/). and NCBI databases and also Wallace Gara for helping with constructing GIS maps

## Data availability statement

All data generated in this study is available upon request from the corresponding author/s

**Table 1:** flow chart of Fig 1

| Number | Geo location | Pango lineage | Samples inside | Diversification events |
| --- | --- | --- | --- | --- |
| 1 | Egypt | B | 52 | 63 |
| 2 | Egypt | B.1 | 15 | 17 |
| 3 | Kenya | B.1 | 89 | 118 |
| 4 | Egypt | AY.116 | 112 | 115 |
| 5 | Morocco | B.1 | 4 | 7 |
| 6 | Ghana | B.1 | 1 | 1 |
| 7 | Ghana | B.1.351 | 4 | 6 |
| 8 | Djibouti | B.1.351 | 100 | 115 |
| 9 | Djibouti | B.1.2 | 23 | 25 |
| 10 | Kenya | B.1.617 | 0 | 1 |
| 11 | Kenya | AY.116 | 0 | 1 |
| 12 | Kenya | AY.116 | 112 | 115 |
| 13 | Kenya | AY.46 | 25 | 24 |
| 14 | Kenya | B.1.617.2 | 5 | 8 |
| 15 | Kenya | AY.16 | 59 | 59 |
| 16 | Kenya | B.1.617.2 | 0 | 1 |
| 17 | Sierra Leone | B.1.617.2 | 23 | 22 |
| 18 | Ghana | B.1.1 | 3 | 9 |
| 19 | Egypt | B.1.1 | 0 | 6 |
| 20 | Sierra Leone | R.1 | 0 | 1 |
| 21 | Sierra Leone | R.1 | 35 | 38 |
| 22 | Egypt | B.1.1 | 13 | 16 |
| 23 | Egypt | B.1.1.1 | 2 | 2 |
| 24 | Egypt | C.17 | 1 | 1 |
| 25 | Egypt | C.36 | 187 | 197 |
| 26 | Egypt | C.17 | 9 | 10 |
| 27 | Egypt | C.36 | 51 | 51 |
| 28 | Egypt | C.36.3 | 94 | 99 |
| 29 | Egypt | None | 3 | 5 |
| 30 | Guinea | None | 0 | 1 |
| 31 | Guinea | Unclassifiable | 6 | 7 |
| 32 | Ghana | B.1.1.7 | 131 | 145 |
| 33 | Djibouti | B.1.1.7 | 0 | 1 |
| 34 | Djibouti | B.1.1.7 | 41 | 70 |
| 35 | Djibouti | B.1.1.7 | 1 | 0 |
| 36 | Zimbabwe | B.1.1 | 5 | 6 |
| 37 | Zimbabwe | B.1.1.111 | 26 | 25 |
| 38 | Ghana | B.1.525 | 0 | 1 |
| 39 | Ghana | B.1.525 | 1 | 2 |
| 40 | Libya | B.1.525 | 27 | 28 |

**Table 2:** Flow chart of Fig 5a

| **Number** | **Geo location** | **Samples inside** | **Diversification events** |
| --- | --- | --- | --- |
| **1** | Djibouti | 2 | 1 |
| **2** | Kenya | 5 | 6 |
| **3** | Malawi | 84 | 222 |
| **4** | Djibouti | 80 | 89 |
| **5** | Djibouti | 2 | 1 |
| **6** | Kenya | 31 | 30 |
| **7** | Seychelles | 18 | 18 |
| **8** | Botswana | 2 | 1 |
| **9** | Kenya/Malawi | 95 | 108 |
| **10** | Djibouti | 24 | 23 |
| **11** | Djibouti | 2 | 1 |
| **12** | Kenya/Malawi | 37 | 38 |
| **13** | Kenya | 44 | 43 |
| **14** | Kenya/Malawi | 24 | 30 |
| **15** | Djibouti | 4 | 3 |
| **16** | Kenya/Malawi | 6 | 9 |
| **17** | Kenya | 7 | 6 |
| **18** | Kenya/Malawi | 4 | 5 |
| **19** | Kenya | 74 | 74 |
| **20** | Kenya/ Malawi | 7 | 7 |
| **21** | Djibouti | 8 | 7 |
| **22** | Kenya/Malawi | 7 | 6 |

**Table 3:**
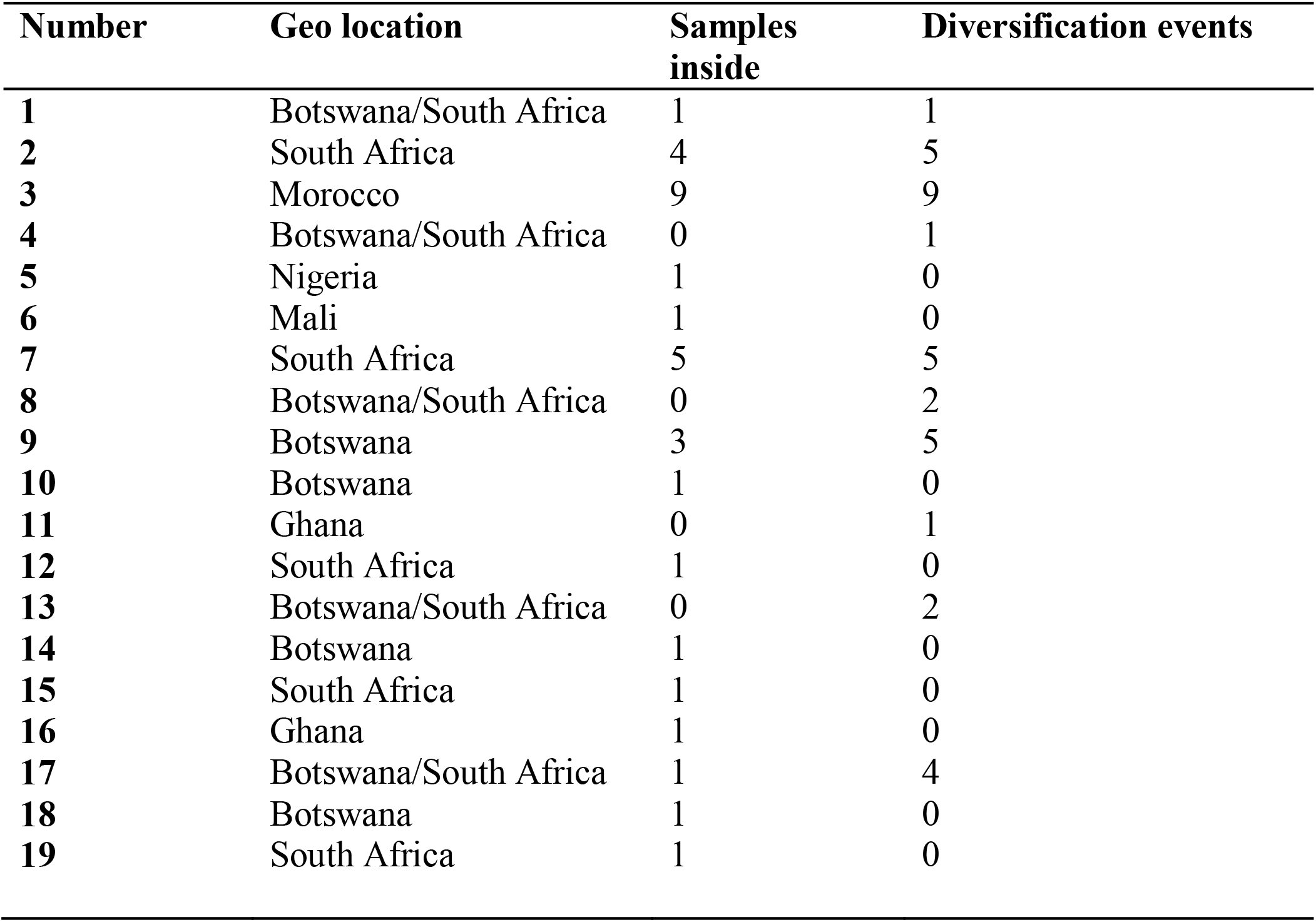
Flow chart of Fig 5b

**Table 4:** Flow chart of Fig 5c

| **Number** | **Geo location** | **Samples inside** | **Diversification events** |
| --- | --- | --- | --- |
| **1** | Nigeria | 91 | 110 |
| **2** | Ghana | 1 | 0 |
| **3** | Nigeria | 1 | 0 |
| **4** | Ghana/Nigeria | 0 | 3 |
| **5** | Kenya | 3 | 2 |
| **6** | Libya | 25 | 24 |
| **7** | Libya | 2 | 1 |
| **8** | Mali | 2 | 1 |
| **9** | Mali | 2 | 2 |
| **10** | Mali | 1 | 0 |
| **12** | Sierra Leone | 1 | 0 |
| **13** | Togo | 1 | 0 |
| **14** | Ghana | 7 | 9 |
| **15** | Ghana | 0 | 1 |
| **16** | Ghana/Togo | 0 | 1 |
| **17** | Ghana | 2 | 1 |
| **18** | Togo | 1 | 0 |
| **19** | Mali | 3 | 2 |
| **20** | Mali | 13 | 16 |
| **21** | Djibouti | 1 | 0 |
| **22** | Ghana | 4 | 3 |
| **23** | Mali | 1 | 0 |
| **24** | Mali | 1 | 0 |
| **25** | Nigeria | 5 | 5 |
| **26** | Ghana | 1 | 0 |
| **27** | Ghana | 1 | 1 |
| **28** | Mali | 1 | 0 |
| **29** | Kenya | 1 | 0 |
| **30** | Nigeria | 1 | 0 |
| **31** | Mali | 1 | 0 |

**Table 5:**
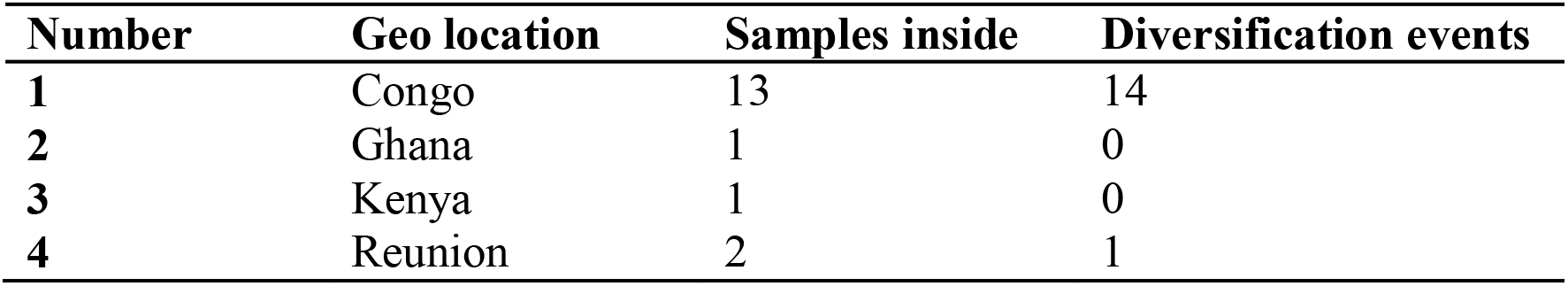
Flow chart of Fig 5d

**Table 6:** Flow chart of Fig 5e

| **Number** | **Geo location** | **Samples**<br>**inside** | **Diversification events** |
| --- | --- | --- | --- |
| **1** | Rwanda | 2 | 2 |
| **2** | Uganda | 130 | 191 |
| **3** | DRC | 2 | 1 |
| **4** | Kenya | 2 | 1 |
| **5** | Kenya/Rwanda/Uganda | 1 | 0 |
| **6** | Kenya/Uganda | 2 | 5 |
| **7** | Kenya/Uganda | 1 | 4 |
| **8** | Kenya/Uganda | 1 | 2 |
| **10** | Rwanda | 35 | 36 |
| **11** | Rwanda | 2 | 1 |
| **12** | Rwanda/Uganda | 1 | 0 |
| **13** | Rwanda/Uganda | 1 | 3 |
| **14** | Rwanda/Uganda | 0 | 1 |
| **15** | South Sudan | 1 | 0 |
| **16** | Zambia | 4 | 6 |
| **17** | Kenya | 5 | 4 |
| **18** | Kenya | 4 | 3 |
| **19** | Kenya/Rwanda/Uganda | 3 | 2 |
| **20** | Kenya | 4 | 3 |
| **21** | Rwanda | 15 | 15 |
| **22** | Kenya/Rwanda/Uganda | 4 | 3 |
| **23** | Uganda | 4 | 3 |

